# The High-throughput WAFFL System for Treating and Monitoring Individual *Drosophila melanogaster* Adults

**DOI:** 10.1101/428037

**Authors:** Maria D.L.A. Jaime, Sean Karott, Ghadi H. Salem, Jonathan Krynitsky, Marcial Garmendia-Cedillos, Sarah Anderson, Susan Harbison, Thomas J. Pohida, Brian Oliver

## Abstract

Non-mammalian model organisms have been essential for our understanding of the mechanisms and control of development, disease, and physiology, but are underutilized in pharmacological phenotypic screening assays due to low throughput compared to cell-based systems. To increase the utility of using *Drosophila melanogaster* in screening, we have designed the whole animal feeding flat (WAFFL), a novel, flexible, and complete system for feeding, monitoring, and assaying flies in a high throughput format. Our system was conceived keeping in mind the use of off-the-shelf, commercial, 96-well consumables and equipment in order to be amenable to experimental needs. Here we provide an overview of the design and 3-D printing manufacture specifications.

## Introduction

Non-mammalian model organisms, such as *Drosophila*, have great potential for phenotypic screens (Willoughby et al. 2013; Strange 2016; Sonoshita and Cagan 2017). They are inexpensive to grow, have a short life cycle, and have well-developed genetic tools for probing biological pathways (Griffin, Binari, and Perrimon 2014; Venken and Bellen 2014; Korona, Koestler, and Russell 2017). These features coupled with the evolutionary conservation of biological processes found in higher organisms like humans have made flies a prime model organism (Adams et al. 2000; Wangler et al. 2017).

Much of high throughput screening, even in flies, relies on tissue culture cells (Esner, Meyenhofer, and Bickle 2018; Heigwer, Port, and Boutros 2018) and does not take full advantage of the model. Metazoans are cells organized into tissues and organs and they exhibit complex systems of interactions among those components. This complexity is why model organisms are so critical for advancing our understanding of development. There is little doubt that the same will be true for pharmacology. While whole organism work captures the important interactions among cell types, tissues and organs, there is an important decrease in throughput. Screens on whole *Drosophila* can easily be scaled to test thousands of conditions, but cell-based screens can scale to millions of samples (Attene-Ramos et al. 2013; Inglese et al. 2007). As a result, *Drosophila* can be used for small scale screening as outstanding follow-up assays following ultra-high throughput screens (Tschapalda et al. 2016; Rimkus and Wassarman 2018) but are not useful as a primary assay at scale. A high-throughput system for screening effects of compounds on flies requires several advances in the way *Drosophila melanogaster* are typically grown in the lab. These advances should be centered on two main aims: first, to reduce the time it takes to manipulate flies within a system amenable to eventual full automation; and second, to reduce the reagent cost of large-scale screens on flies.

*Drosophila* are typically grown in vials with 10 mL of food, but they can also be grown in deep-well 96-well plates using < 1 mL of food (Willoughby et al. 2013; Markstein et al. 2014). However, this is still a large volume if one is interested in screening small molecule libraries, where the amount of any given compound is low and as a result miniaturization greatly augments scale (Kenny et al. 2015). Additionally, while simple fly assays such as viability are relatively easy to scale, it is difficult to perform a range of behavioral and biochemical assays without extensive retooling and protocol development.

One of the biggest problems is that treated flies, the food they consume, and excrement they produce are all in the same well. It is possible to develop elegant ways to separate flies from the media, such as floating larvae to the surface for imaging (Willoughby et al. 2013), but a simple and flexible method to recover the flies following treatment, along with measurement of food intake and harvest of excreted material, would greatly improve throughput and the range of assays that could be performed on treated flies. There are methods by which small volumes of food are supplied externally to where the flies are housed, but these are low throughput. The Capillary feeder (CAFE) assay delivers microliters of liquid food to flies using capillary tubes inserted in the lids of narrow vials (Ja et al. 2007). The Expresso assay (Yapici et al. 2016) is a modification of the CAFE assay with multiple single-fly feeding chambers, each connected to a sensor bank that delivers and measures the amount of liquid food consumed by sets of individual flies. These liquid feeding methods provide excellent measurements of media consumption and the flies can be harvested without bringing the food into the assay, but they do not scale.

The Whole Animal Feeding FLat (WAFFL) uses the advantages of deep-well plates and CAFE-like systems, while being scalable and versatile in form and function. We kept several design criteria in mind as we developed WAFFL prototypes. To boost throughput, including compatibility with eventual robotic screening, and to avoid having to develop excessive numbers of custom components, we sought compatibility with commercially available consumables and equipment, such as 96-well plates, 96-tip pipette heads, 96-well silicone mats, 96-well plate centrifuge rotors, and 96-well homogenizers. We were interested in minimizing manipulation of the flies to prevent escape and disturbances to flies in potentially sensitive assays. This included the ability to easily harvest flies for assays while comprehensively tracking each individual fly treatment. We additionally sought to use small volumes of media in order to support future experiments where cost or availability of reagents is a driving factor in feasibility. The 3D-printed WAFFL system uses a set of interlocking components to allow for measurements of consumption, behavior, and biochemical assays with essentially no manual intervention after the flies are loaded into the system. The WAFFL system is easy to use and compatible with a broad range of assays.

## Results

### System Overview

We developed the 96-well Whole Animal Feeding FLat (WAFFL) system to rapidly perform high-throughput experiments and deliver essentially any compound or diet in 16 microliters of liquid media, enough for 18 hours of exposure. We designed a housing plate, in which flies feed through a series of pores at the bottom of each chamber when the housing plate is placed into a standard 96-well microtiter round bottom plate with liquid food (Figure 1A, S1A). Because we designed the housing and liquid food plates as separate components, the media can be changed as desired without manipulating the flies (Figure 1B). One simply prepares a new 96-well plate of food, starvation media, or other test substance, and the housing unit is moved from the old to the new feeding plate. This was designed to allow for a variety of experimental designs, such as the effects of order of delivery to the flies, or the effects of an acute bolus versus chronic exposure.

**Figure 1.**
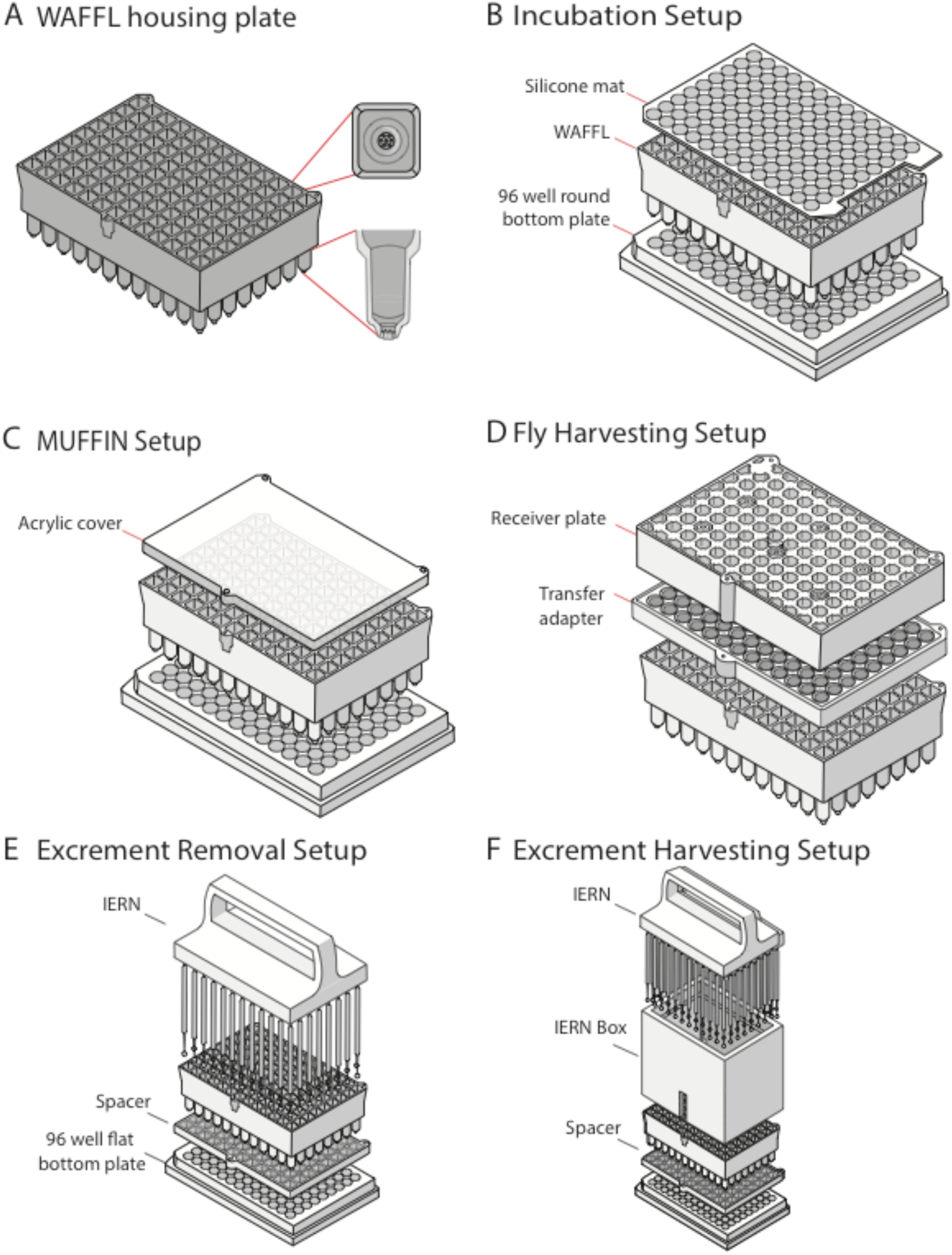
WAFFL diagrams and setups. A) 3-Dimensional representation of the WAFFL housing plate as seen from above. Two insets show a top and side view of an individual chamber. B) The incubation setup used a WAFFL housing plate with flies, a silicone mat seal, and a 96-well plate with liquid food. C) The MUFFIN filming setup was like (B) but a clear acrylic cover replaces the silicon mat. D) The fly harvesting setup used the transfer adapter and a receiver plate to move the flies from the WAFFL to the 96 deep-well plate in the same well orientation. We flipped this setup upside down to harvest flies and retransferred to a 96 deep-well plate for further processing (not shown). E) Step one of the excrement removal setup used the WAFFL-IERN with a spacer to remove the excrement from the wells of the WAFFL housing plate. F) Step two of the excrement setup used the IERN box to provide support to the WAFFL-IERN while the whole device was centrifugated to collect the excrement in the 96-well flat-bottomed plate.

The WAFFL housing chambers have square cross sections at the top (to maximize living quarters for the flies) and circular cross sections at the bottom (for fitting into standard plates). The top part of the chambers are rectangular prisms, with the long axis in the Z-plane (8 mm X 8 mm X 24.5 mm) (Figure 1A, S1A). At the base of the rectangular portion, there is an inner edge that protrudes 0.4 mm from the inner walls (Figure 1A). Each chamber has a conical protrusion (11.3 mm in length) with a slightly flatted bottom and seven evenly spaced 350 μm diameter holes (Figure 1A, S1A). The volume of each chamber is 1.6 mL. The housing plate also includes nine asymmetrically located orifices where steel alloy dowels are fitted and then sealed with clear silicone to allow the housing plate to magnetically attach to the complementary pieces of the transfer system when the pieces are matched in the correct orientation. This design reduces the chances of mislabeling samples due to plate rotations during the course of an experiment.

The top of the housing plate is compatible with 96-well plate silicone mats (Figure S1B) and custom acrylic lids (held by dowels) for video monitoring (Figure S1C), using the Monitoring Unit for Fruit Fly Imaging in Ninety-six-wells (MUFFIN, Figure 1B, 1E, 2). This system allows us to see if the flies are interacting with or avoiding the media, and for scoring behavior on treatment.

Flies can be harvested at the end of an assay by the transfer system, which recovers flies for essentially any assay, such as microscopy or homogenization in 96-well format with stainless steel beads (Figure 1C, S1E - F). Many of our assays involve homogenization, and beads can interfere with pipetting the supernatant, so we developed a 96-magnet bead remover (Figure S2D). Because the food and housing chambers are distinct, we can also harvest the fly excrement at the end of an experiment using the WAFFL Insect Excrement Removal Nano-brushes (WAFFL IERN) component of the system (Figure 1D, S2A - C).

A short video showing the use of the WAFFL is available (YouTube https://youtu.be/6yJ4Mq8xIXc and supplementary file).

### WAFFL housing unit

The housing plate is the main element of the WAFFL system (Figure 1A, S1A). Up to three flies can inhabit each of the 96 chambers in the WAFFL, allowing for some pooling to dampen individual variance, but the real strength is the ability to measure those individual differences by placing a single fly per chamber. In the production model, each chamber has a volume of 1.6 mL, which provides ample room for movement (Figure 1A - B). Previous work has shown that 1.1 - 1.3 mL 96-well plate formats with solid food are effective for screening flies (Willoughby et al. 2013; Markstein et al. 2014), so the WAFFL housing chamber is spacious. The upper portions of the WAFFL plates are square, to provide maximum volume per well while also being compatible with commercially available silicone mats that can be used to seal flies within the WAFFL plates (Figure 1A - B, S1B).

The WAFFL is designed to use liquid food. The food access points at the bottom of the WAFFL are critical and were one of the most challenging parts to design (Figure 1A). We experimented with several different designs focused on the spacing between the WAFFL housing unit, the inside of commercial 96-well plates, and the access ports that allowed the flies to feed. To make a tight fit, we implemented a design with square chambers on the upper portion of the WAFFL that transitioned into a cupular projection at the bottom of the housing unit. This projection fits inside the round well plate with 1–2 mm clearance between the wall of the WAFFL housing plate and the well of the food plate. Tighter clearance distances did not provide enough room for liquid in the food plate and there was excessive capillary action (this also depends on the viscosity of the media) between the WAFFL and the food plate, including the spaces between wells. Capillary action exterior to the housing chamber did not allow adequate food access for the flies and resulted in high mortality. More loosely fitting plates constrained flies at the feeding position and increased the volume of media used in each well.

We tested different types of openings for the cupular feeding ports where the media is accessible to the fly. This included proboscis-sized sipping ports and larger openings, giving flies access to a media pool. We observed that when we provided a large opening or many openings at the bottom of the WAFFL plate, flies often became trapped in the liquid media. This did not occur when we reduced the size of the openings and provided a horizontal surface near the feeding port (at the transition from the square to conical shape) where flies could sit. This perch also helped keep flies that fell while wall climbing from falling directly into the feeding area.

Small ports cannot be 3D printed by low-resolution additive manufacturing devices. For our final design, we determined that a high layer resolution of 0.05 mm (Z) and in plane resolution of 0.08 mm (X - Y) was sufficient. In the production model, we created seven 350 *μ*m diameter openings at the slightly flattened terminus of the cupular projections. This arrangement and size provided a good balance of space for the flies to feed, without falling due to the presence of excess liquid.

When considering the material used for printing the WAFFL, we needed to ensure that the housing plate was waterproof so that the liquid in the food plate that we place the WAFFL into was not absorbed by the plate material and could be easily cleaned. The material also needed to be relatively non-reactive with the compounds we might use in assays, biosafe, structurally stable, and not bleed during the printing process as to not alter the specified dimensions. After evaluating and testing the ability of the materials to meet our requirements, we chose Accura ClearVue, a USP (United States Pharmacopeia and National Formulary (USP - NF) plastic designation) class VI capable, transparent, and bio-compatible stereolithography (SLA) resin.

We have tested loading flies into the WAFFL as 2^nd^ or 3^rd^ instar larvae or pupae (1^st^ instar larvae can pass through the feeding ports) but, in most work, we load adult flies. We printed a custom four-channel fly dispenser (Figure S1D), which aspirates (by mouth) four flies at a time for transfer into the housing plate. To keep the flies anesthetized during loading, we designed and printed a WAFFL CO_2_ base (Figure S2E). The housing plate is securely seated in the CO_2_ base and humidified CO_2_ enters through the seven openings at the bottom of the cupular section of each well. These additional components facilitate faster and more consistent fly loading.

The assembled WAFFL housing plate and food plate should be wrapped with parafilm to minimize evaporation during an experiment longer than 8 h. Additionally, we use plastic boxes to hold groups of plates in a humidified environment. The plates sit in a base above a saturated salt solution selected to maintain a desired (high) humidity level in the chamber (Figure S2F). Groups of plastic boxes are then placed in a larger incubator.

### Video monitoring

There are many fly behavior monitoring systems (Simon and Dickinson 2010; Gilestro 2012; Kohlhoff et al. 2011), however these are not designed to perform an assay with 96 flies and simultaneously record high resolution video, while also being easily portable and compatible with other 96-well equipment. Therefore, we developed a new video system for the WAFFL.

The Monitoring Unit for Fruit Fly Imaging in Ninety-six well (MUFFIN) was designed to capture high resolution (720p) and frame rate (30 fps) video for each chamber in the WAFFL to track fly position and enable detection of certain behaviors of interest (Figure 2). Aside from the image resolution requirement, practical constraints for the MUFFIN design were mainly compactness and cost-effectiveness. The MUFFIN employs 24 Raspberry-Pi cameras fitted with standard lenses, each of which was focused on a grid of four wells (Figure 2E, 2I). This number of cameras allows capture of the field of view with sufficient resolution in a small working distance, allowing for a compact system.

**Figure 2.**
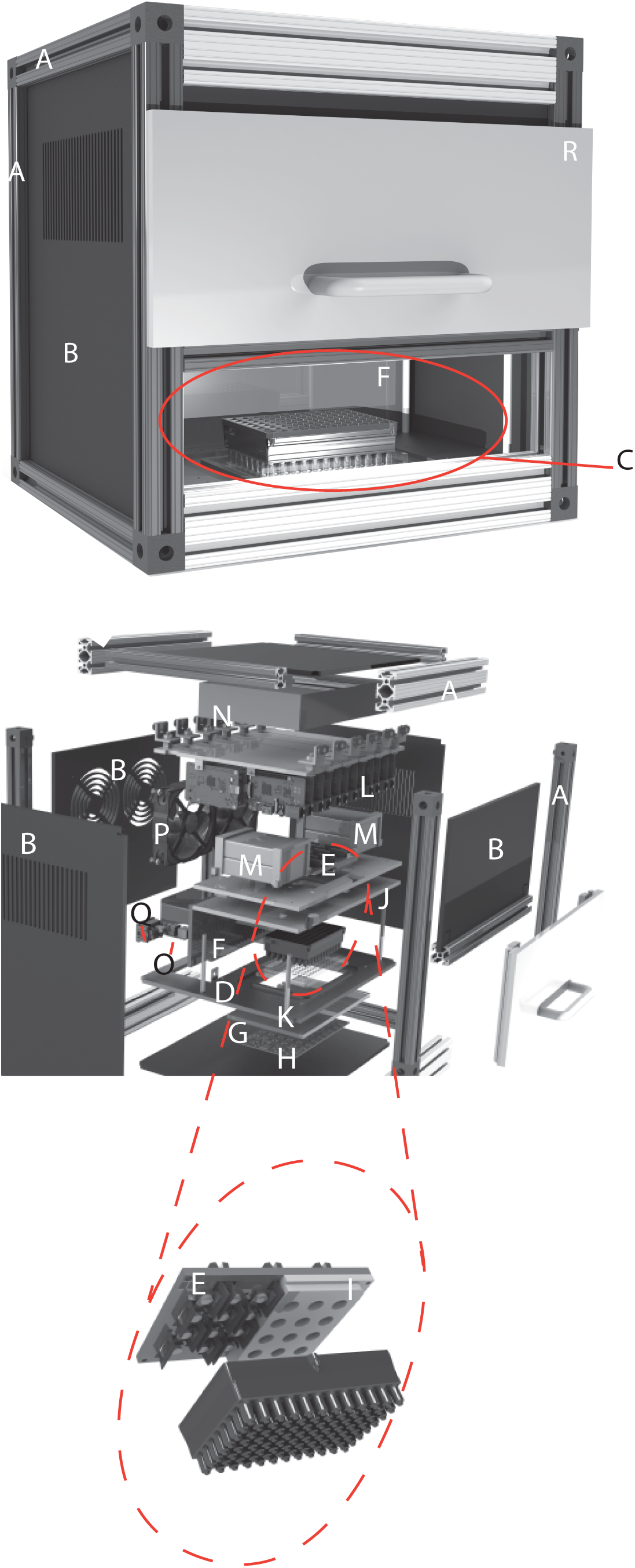
Exploded diagram of the Monitoring Unit for Fruit Fly Imaging in Ninety-six wells (MUFFIN). A) slotted aluminium frame. B) acrylic covers. C) monitoring chamber. D) acrylic base plate. E) camera sensors and lenses. F) acrylic wall. G) mounting cutout with near-infrared (NIR) light (700 nm) diffuser. H) printed circuit board. I) 3D printed camera board mount. J) acrylic support. K) support stand-offs. L) 24 Raspberry Pi (RPI) modules. M) USB power ports for RPIs. N) 24-port Ethernet switch and a five-port Ethernet switch. O) output cable. P) fans. Q) power entry module. R) sliding lift door.

**Figure 3.**
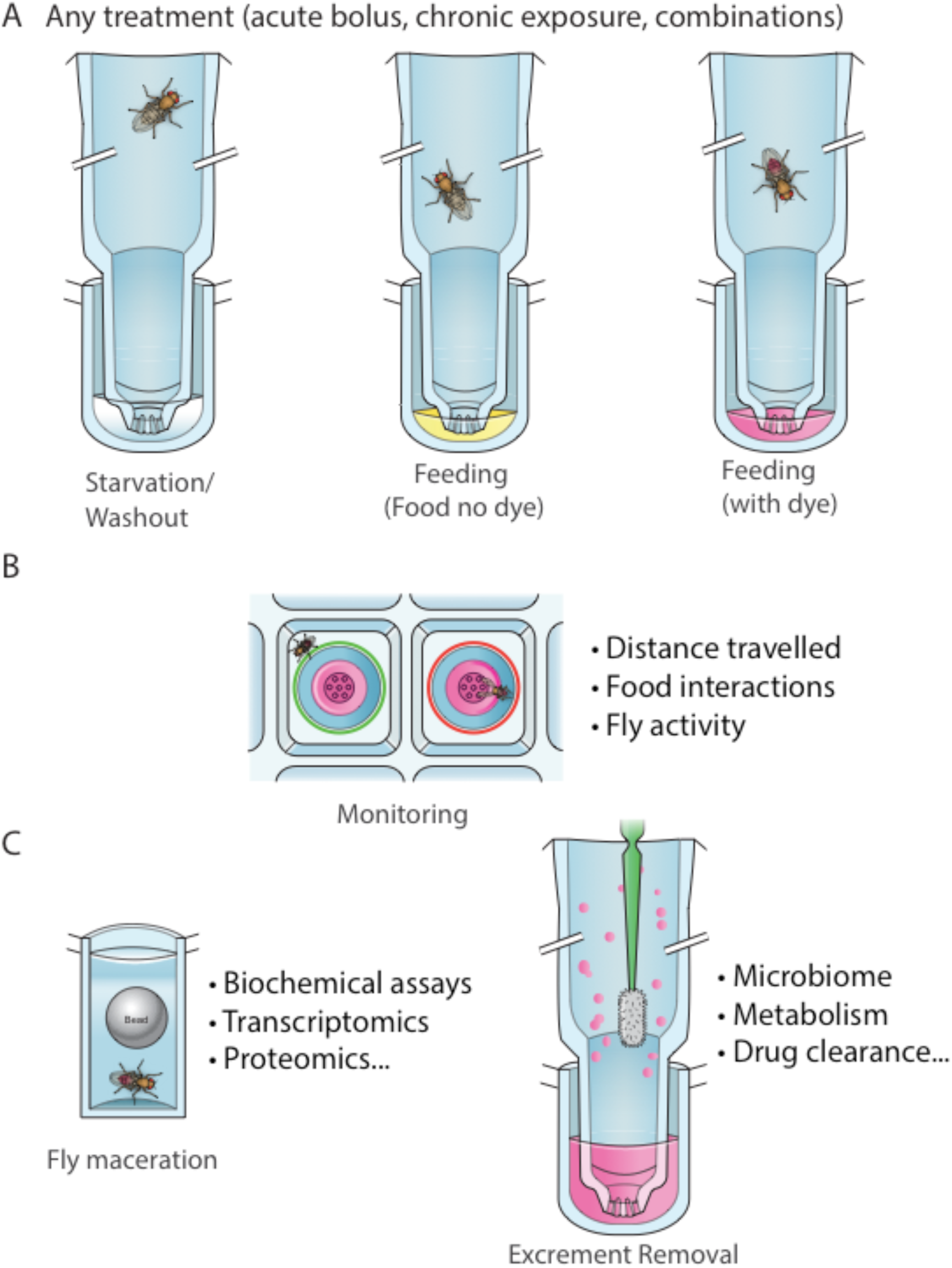
Overview of the WAFFL system capabilities. A) One fly is placed in each well of the housing plate. Example treatments of starvation, feeding, and feeding with tracking dye. B) Example of MUFFIN monitoring for recording proximity to food. Top view shows a fly outside the region of interest (red circle, left) and within the area of interest (green circle, right). C) Fly collected for homogenization assays (left) or excrement harvesting (right).

To avoid compromising the results of MUFFIN experiments with confounding factors introduced by the system components (such as heat and noise), and to protect the electronics from the corrosive volatile hydrocarbons that flies produce, the MUFFIN was built with a recording chamber physically isolated from the rest of the system’s electronics and components (Figure 2F). The recording chamber was also separated from the external environment with a pull-down door (Figure 2F). The recording arena has a keyed plate to hold the WAFFL directly underneath the system cameras, such that each camera’s field of view was fixed and consistent between recording sessions (Figure 2C). Each of the 24 Raspberry Pi’s streamed video to a local computer via a local gigabit network. We saved video on a computer and watched the progress of the live recording by streaming.

For experiments in the MUFFIN, we placed a thin clear acrylic plate cover on the WAFFL to prevent flies from escaping without obstructing the cameras view (Figure 1E, S1C). We tested several clear acrylic covers of different thicknesses varying from 1.59 mm to 6.35 mm, to choose the one resulting in the highest image quality. Thicker acrylic pieces resulted in darker images, and thinner acrylic resulted in distortion due to flexing.

We placed the MUFFIN inside the same incubator as the WAFFL experiments to make the conditions consistent. In our current work, the conditions were an average temperature of 25 ^o^C and average humidity of 75 %. The temperature inside the MUFFIN rose up to 27 ^o^C as result of the electric parts of the device. We chose a high humidity to avoid evaporation of the liquid media due to its low volume of 16 *μ*L. Having a higher humidity also prevented condensation of the liquid (probably from evaporating food) on the acrylic cover.

The chosen MUFFIN cameras were versatile in recording resolution and acquisition rate. We have developed a first generation set of custom algorithms that detect fly silhouettes in the image and identify the centroid of the silhouette of the fly. The silhouette can be used to compute a measure of the frequency of interactions with food, while the centroid is used to compute a measure of the locomotive activity of the fly in terms of image distance traveled in the X - Y axis. Although the software does not provide estimates for Z axis position, measurement of movement in the Z plane should be possible based on statistical learning methods (Salem et al. 2015).

### Harvesting Flies

We wanted to design a method to transfer the flies from the housing plate to a deep-well plate for tissue homogenization and further assays. This can be simply achieved by inverting a fresh plate, inverting the paired plates, and tapping the flies into the new plate. However, to facilitate eventual robotic implementations, we were interested in transfer to a range of plate types and also wanted to promote better tracking by demanding a single plate orientation. Therefore, we decided to add a series of possible intermediate adaptors that retain the orientation of the plate throughout the experiment and permit flexibility (Figure 1C). As a tracking feature, we added an asymmetrical orientation locking system via 9 pairs of steel dowels and magnets in each piece of the system, ensuring a unique alignment of the three components of the WAFFL system (housing, receiver plates, and transfer adapter). This eliminated the possibility of rotating plate orientation during processing and subsequently mislabeling the samples. Anesthetized or frozen flies can be flipped and centrifuged to transfer.

For many experiments, such as RNA-seq library construction, reporter expression, or metabolite measurement, homogenization is required. In order to macerate the flies, we simply added a stainless-steel bead into each well of the deep 96-well plates containing the flies with the selected buffer for use and loaded the assembly into a bead beater. Following homogenization, removal of supernatant with a pipette from the deep well plate with the 5 mm stainless steel bead inside each well was difficult and inaccurate, as beads block the pipette tips. To address this, we designed and printed a 96-pronged magnetic device to remove the stainless-steel beads (Figure S2D). The key design challenge was that each of the 96 prongs needed to have a magnet strong enough to grab the bead, but not so strong that the magnets would interact with each other and deform the prong geometry. We chose to encapsulate two nickel plated axially magnetized Neodymium magnets (3.18 mm diameter X 6.35 mm thick) in each prong. As configured, the bead remover was able to remove each bead in one swift motion, making the processing of the homogenized fly tissue more efficient.

### Harvesting excrement

Most quantification of media consumption relies on measuring residual non-consumed media. We decided to actively measure the amount of media that passed through each fly by adding non-metabolizable, non-absorbable tracer dyes to the media and measuring excrement (Ja et al. 2007). Measuring excrement has the added advantage of making it possible to assay for a range of poorly studied parameters, such as metabolites, drug clearance, and effects on the microbiome. The WAFFL Insect Excrement Removal Nano-brushes (WAFFL-IERN) was designed to aid us in removing the feces of the flies that underwent screening in the WAFFL (Figure 1D, S2A-C).

The main part of the WAFFL-IERN (Figure 1D, S2A) is a handle with 96 micro brushes that align with the WAFFL housing chamber plate. After the food plate and flies are removed, the WAFFL housing plate with excrement is placed into a 96-well plate with a suitable buffer (we have used PBS pH 7.4 + 0.5% v/v Triton) and the WAFFL-IERN is used to manually brush the interior of the WAFFL chambers with a series of up and down movements. The WAFFL-IERN fits into a housing that supports the handle, such that the brushes are unable to reach the bottom of the WAFFL. The WAFFL-IERN and housing is centrifuged to remove liquefied excrement for collection in a 96-well plate (Figure S2C). We designed a spacer for use during centrifugation to raise the WAFFL 5 mm from the base of the flat bottom 96-well plate so that the excrement solution does not touch the tip of the cupular wells of the WAFFL (Figure 1D, S2B).

We have provided a detailed description of the WAFFL and its complementary components, the MUFFIN and the WAFFL-IERN. In conjunction, these devices provide a comprehensive system that makes performance of high-throughput screens, evaluation of excreta, and monitoring of adult flies plausible.

## Discussion

The WAFFL system is a unique set of designed and 3D printed labware that uses standard 96-well equipment to support a wide-range of predicted use cases, rather than a particular assay. This system minimizes the amount of food or media and fly manipulation required and was designed to be easy to automate. The WAFFL system fills a gap in the set of non-mammalian model organism screening systems currently available. The system is currently made by additive manufacture, which has distinct advantages for customization by users for other uses or model organisms. The system could also be injection-molded to further reduce per unit cost.

The WAFFL system allows performance of a series of assays with the same set of flies. Fly activity and food interactions can be monitored by the MUFFIN, indicating the effects of a treatment on fly behavior. Having the flexibility to change the food within seconds allows studies aimed at understanding the effects of acute bolus versus chronic exposures as well as determining the effects of order of delivery to the flies.

Excrement can be harvested after treatment with the WAFFL-IERN to evaluate metabolism, drug clearance, and treatment effect in the microbiome. Treated flies can be easily collected after treatment and evaluated individually by microscopy or tissue homogenization for assays such as transcriptomics, proteomics, and metabolite measurements. The combination of assays increases the number of measurements per fly, leading to better understanding of the biological question. Using manual and semi-automated protocols, we have performed up to 20,000 fly experiments in a week. The only serious impediment to screening we observed was evaporation of liquid media when performing long assays (>8 h) or when a heat shock or elevated temperature is part of the experimental design. This is a common problem in essentially all high-throughput assays on cells (Maddox, Rasmussen, and White 2008). However, a major advantage of the WAFFL is that since flies are kept in the housing plate one can simply replace the media plates as needed for longer assays.

While the one-fly one-treatment per well design is a strength in many settings, there may be situations in which a researcher wants to co-house flies or offer flies a choice of media. Well size could be increased by modifying our current design. For example, merging two or four top square sections of the well so the well size doubles or quadruples would permit housing more flies per well. In this larger format (4 merged wells), if there were four conical projections to the feeding plate, one could provide flies the option of any of four different media that could be mixed with different dyes to assay food preferences. One might also want to reduce the z dimension to create an arena rather than a deep well. This could be especially useful for complex behavior assays.

In summary, we developed a comprehensive system that could support both parallel and sequential assays in adult flies using the same platform. We envision that the WAFFL system has the potential to revolutionize the use of phenotypic screening in Drosophila and inspire the development of related systems for other small model organisms.

## Materials and Methods

The Author Reagent Table (Table 1) lists all of the materials and reagents used to produce the various apparatus. Design associated print files and software for described components can be found in the supplement for use by academic researchers. The WAFFL design files for printing the system described herein are available under a material transfer agreement (MTA) from the Technology Advancement Office (TAO) of the National Institute of Diabetes and Digestive and Kidney Diseases (NIDDK), an institute of the National Institutes of Health (NIH). Interested parties should contact. The WAFFL system is available for commercial license from TAO-NIDDK. Contact for licensing inquiries.

### WAFFL components

The WAFFL housing plate and adaptors were designed with Autocad 2014 for Mac (Autodesk, San Rafael CA). We generated the data file using 3D Lightyear 1.5.2 and sent the file to the 3D printer with Build Station 5.5.1 (3D Systems, Rock Hill SC). For the WAFFL CO_2_ base, Four Channel Fly Dispenser, bead remover and WAFFL-IERN we used SolidWorks 2018 (Dassault Systems, Waltham MA) and Insight 9.1 to create the toolpath/data file (.CMB file format) and Control Center 9.1 to send the .CMB file to the 3D printer (Stratasys, Eden Prairie MN).

The WAFFL housing, adaptor and transfer plates were made by stereolithography printer in high resolution mode (X and Y = 0.08 mm, Z = 0.05 mm) on a Viper 2Si SLA (3D Systems, Rock Hill SC) from Accura ClearVue resin (3D Systems, Rock Hill SC). Pieces were post-processed in a glycol/ether bath for 2 h to soften the supports and remove resin residues, soaked in isopropanol for 30 min, and cleaned with a toothbrush and dental files (25 mm size 030). Pieces were dried with house compressed air and, in the case of the WAFFL housing plate, we used compressed air to check the seven openings at the bottom of each chamber. We used an ultrasonic bath (Quantrex, Kearny NJ) with isopropanol to remove any leftover resin. Pieces were air dried and then UV cured in a ProCure 350 oven (3D Systems, Rock Hill SC) for 40 min. The cross sectionally square end of the chambers was filed with a square needle 2.7 mm file (Harbor Freight Tools, Camarillo CA).

The WAFFL CO_2_ base was made on a PolyJet printer (Eden260VS, 16-micron layers, X/Y-axis: 600dpi) with VeroClear plastic (Stratasys, Eden Prairie MN) with a glossy surface finish. The support material (SUP707) was removed in water using a circulating SRS-DT3 tank (CleanStation, Osseo MN) and air dried. We drilled a hole for CO_2_ input to the hose barb fitting (19.05 mm), which was attached with a locknut fitted with Teflon tape.

The fly dispensers were printed on a PolyJet printer (as above), with Rigur plastic (Stratasys, Eden Prairie MN) and a matte surface finish. Support material (FullCure SUP705) was removed using an OBJ-01200 water jet (Stratasys, Eden Prairie MN) and 1 mm metal rods were used to completely clear the holes, followed by air drying. A rubber tubing (1.95 mm) was attached to aspirate the flies.

The bases and spacers for the WAFFL-IERN humidity chambers and other simple pieces were made from cast acrylic cut with a PLS6.150D laser (Universal Laser Systems, Scottsdale AZ). We annealed the liner pieces using a 40AFE-LT, 800W lab oven (Quincy Lab Corporation, Chicago IL) at 80 ^o^C to prevent cracking.

The bead remover, WAFFL-IERN handle, and IERN support box were printed by fused deposition modeling with ABSplus plastic using the Fortus 250mc printer (Stratasys, Eden Prairie MN). The support material was removed in the SRS-DT3 tank (CleanStation, Osseo MN) filled with 41.6 L water and 950g NaOH, rinsed in water and air dried. Each of the prongs of the bead remover had two thick nickel plated axially magnetized Neodymium magnets (Apex Magnets, Petersburg, WV) of 3.18 mm diameter X 6.35 mm inserted and sealed with Loctite M-31CL epoxy (Henkel Corp, Westlake OH). Disposable 1.5 mm micro-applicators were fitted manually (Microbrush, Grafton WI) into the handle of the WAFFL-IERN. The micro-applicators can be replaced as needed.

### Video monitor

The MUFFIN enclosure is composed of a slotted aluminum frame (80/20 Inc., Columbia City, IN), with 3.18 mm thick black cast acrylic covers (Piedmont Plastics, Charlotte, NC) mounted on the frame. The unit has a monitoring chamber for the loaded WAFFL. The remainder of the unit houses the electronic and optical components.

The monitoring arena was assembled from the following: an acrylic base-plate for securing the WAFFL, a near-infrared back-lighting subsystem placed directly beneath the base-plate, an array of camera sensors/lenses mounted above the base-plate, and an acrylic partition to isolate the monitoring arena from the other components.

The base-plate had a cutout matched to the dimensions of the well-plate (2mm clearance). A white translucent acrylic near-infrared diffuser (Piedmont Plastics, Elkridge, MD) mounted on the bottom side of the base-plate supported the WAFFL in the cut-out. The acrylic mounted 2.54 cm beneath the printed circuit board provided near-infrared light. The custom designed board consisted of an array of LED emitters (Vishay Semiconductors, Malvern, PA) and associated driver circuitry (Linear Technology, Milpitas, CA). The combination of the LED array and diffuser achieved spatially uniform backlighting for the WAFFL. Above the WAFFL was an array (Figure 2E) of 24 Raspberry Pi NoIR camera boards (Raspberry Pi Foundation, Cambridge, UK) placed in a 6 x 4 grid such that each lens field-of-view was a 2 x 2 well region within the 96-well WAFFL format. The camera boards incorporate the sensor/lens assembly, electronically connected via a flexible PCB cable and mechanically mounted at the center of the board via an adhesive pad. The physical dimensions of each camera board exceed the dimensions of the 2 x 2 well region within a 96-well plate. To achieve the desired positioning of the lenses, the adhesive was removed to free the lens, allowing it to be repositioned relative to the camera board on a custom 3D-printed part (partially hidden in the figure to reveal underlying detail of camera board and lens placement). The lens was designed with slots to interleave and secure the camera boards in press-fit cutouts, such that each lens has a perpendicular top-down view of its assigned 2 x 2 region. This lens/camera board mounting part is held from above by an acrylic plate suspended above the base-plate by stand-offs (McMaster-Carr Supply Co, Elmhurst, IL). Lastly, to isolate the monitoring arena from the other components in the system, an acrylic sheet was mounted perpendicularly between the base-plate and the camera mount acrylic.

In addition to the monitoring arena, the MUFFIN unit housed the electronics needed for the back-lighting subsystem, as well as the video acquisition and transmission: a 12V power transformer (Qualtek Electronics Corporation, London, ON) to supply the illumination PCB, 24 Raspberry Pi (RPI) modules which acquired video from the camera boards, USB power ports (Anker, Seattle, WA) to supply the RPIs, a 24-port Ethernet switch (TRENDnet, Torrance, CA) and a five-port Ethernet switch (Netgear, San Jose, CA) used to transmit the video data from all 24 cameras to a single destination (e.g., an external control computer) via an Ethernet cable. The sole output cable from the Ethernet switch was connected to a panel-mount Ethernet feed-through connector (Amphenol Corp, Wallingford, CT). Fans (MITXPC, Fremont, CA) are mounted on the back-cover for system thermal management. A power strip with seven receptacles (Tripp Lite, Chicago, IL) was installed for use by all devices requiring AC power. The main power cord of the power strip was stripped and soldered on to a power entry module (TE Connectivity Ltd, Schaffhausen, Switzerland) mounted on the back cover of the unit.

The rear cover of the MUFFIN enclosure had a power entry module for a standard AC power cord and an Ethernet cable receptacle. The front of the MUFFIN enclosure had a sliding lift door (Figure 2R) allowing user access and isolation of the monitoring arena.

### Video software

Custom software ran on both the Raspberry Pi boards and the external control PC in order to stream the videos over a MUFFIN-limited Ethernet LAN for storage on the PC. The Python-based MUFFIN acquisition software, with a graphic user interface (GUI), recorded video from the 24 Raspberry Pi cameras enclosed in the system (Figure S4). The software interfaces with the Raspberry Pis over a local gigabit network. The GUI consisted of a video display window surrounded by status indicators and control buttons. The display window was divided into four sub-windows, allowing the user to view live video from four separate groups of wells at one time. Status indicators and controls are located along the right side and bottom of the GUI. The uppermost group of status indicators on the right side showed experiment information (name, start time, end time, and total runtime). Below, a section for computer information (Figure S4C) included hard drive space, host PC IP address, and number of connected cameras. The well selection area window (Figure S4D) contained four arrays of checkboxes to toggle the groups of wells displayed in the video display windows. This section also contained a button to manually start and stop recording (Figure S4E). Finally, the bottom of the GUI contained a group of buttons that controlled the program settings and displayed the current date and time (Figure S4F). From left to right, this section included buttons to close the application, minimize the application, display time and date, add experiment information, and adjust program settings.

The minimum computer requirements for the MUFFIN video acquisition and GUI in our hands were an Intel Core i5-6xxx or equivalent, Microsoft Windows 7, 8 GB RAM, 500 GB internal hard drive, Gigabit wired ethernet port, and USB 3.0 ports.

## Author contribution

BO and MDLAJ conceived the idea of WAFFL. MDLAJ designed the WAFFL system. MDLAJ, GHS, JK, MG, SH, TJP conceived the idea of the MUFFIN. GHS, JK, MG, SA, TJP designed and assembled the MUFFIN. GHS, JK and TJP developed the MUFFIN algorithm. SH provided advice and materials for the MUFFIN. BO, SK and MDLAJ wrote the manuscript. SK and MDLAJ performed validation assays with the MUFFIN. All authors read and approved the final manuscript.

## Acknowledgments

We are grateful to Dr. Gerald T. Grant, Dr. John P. Lichtenberger and Dr. Peter Liacouras from the 3D Medical Application Center at Walter Reed National Military Medical Center (3D MAC WRNMMC) for technical advice and allowing us to use their 3D printers to make possible the prototyping of the WAFFL systems. Thank you to Cale Whitworth and Hangnoh Lee for advice and discussions in different aspects of this project. This research was supported by the Intramural Research Program of the NIH, in the National Institute of Diabetes and Digestive and Kidney Diseases (NIDDK) and the National Heart Lung and Blood Institute (NHLBI).

**Abbreviations**
GUI: :Graphic User Interface
IERN: :Insect Excrement Removal Nano-brushes
LED: :Light-Emitting Diode
MTAs: :Material Transfer Agreements
MUFFIN: :Monitoring Unit for Fruit Fly Imaging in Ninety-six-wells
NIDDK: :National Institute of Diabetes and Digestive and Kidney Diseases
NIH: :National Institute of Health
NIR: :Near-Infrared
PCB: :Printed Circuit Board
SLA: :Stereolithography
WAFFL: :Whole Animal Feeding FLat
3D: :Three-dimensionality

## Supplementary figures

**Figure S1.**
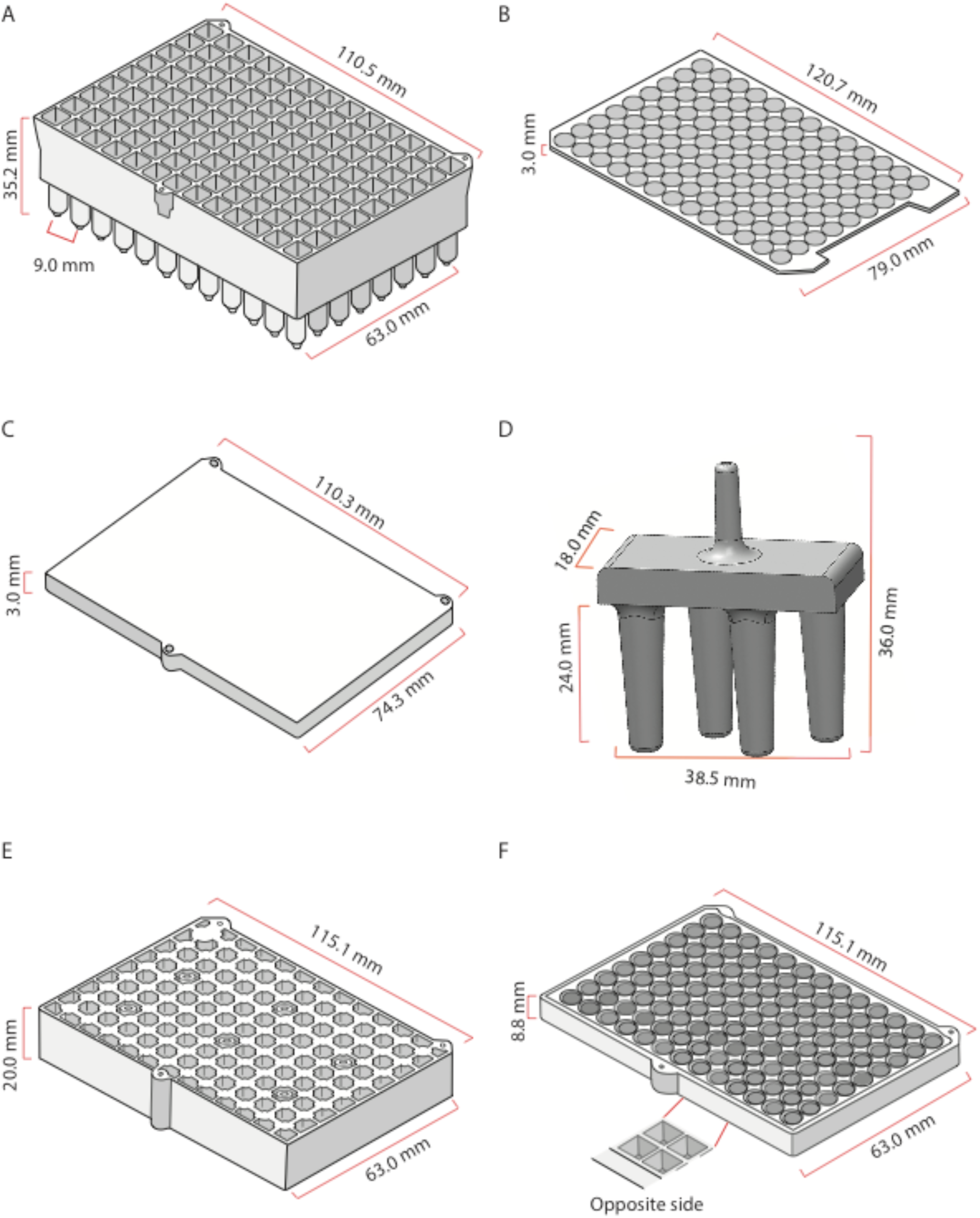
This figure provides more detailed measurements for each of the pieces of the WAFFL system. A) Housing plate, B) Silicone cover, C) Clear cast acrylic cover for MUFFIN assays, D) 4 channel fly dispenser, E) Receiver plate, F) Transfer adapter.

**Figure S2.**
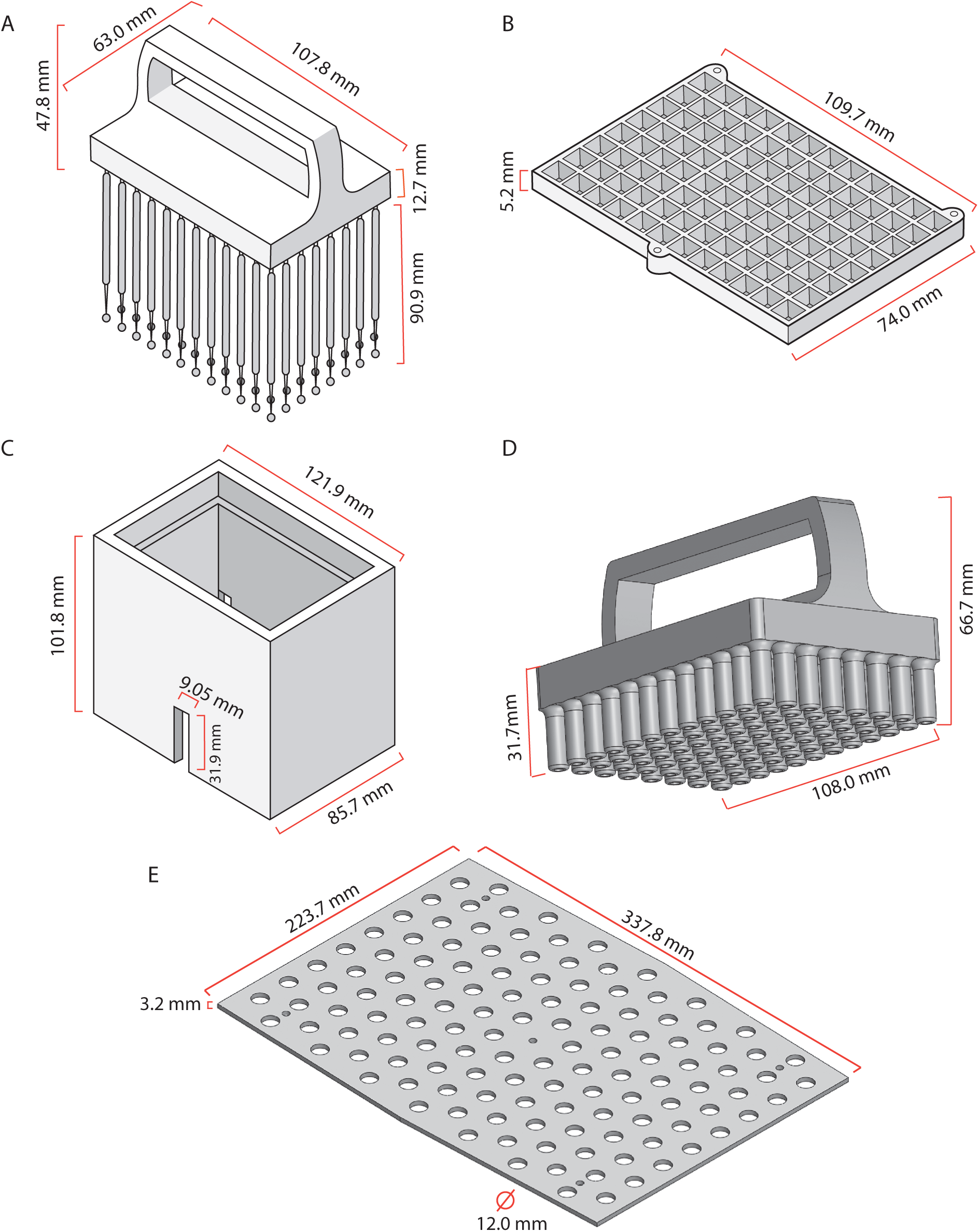
This figure provides more detailed measurements for each of the accessories for the WAFFL. A) Insect Excrement Removal Nano-brushes (WAFFL-IERN), B) 96 well spacer, C) IERN Box, D) 96-well stainless-steel bead remover, E) Rectangular base-board for stabilizing the WAFFL in the incubation chambers.

**Figure S3.**
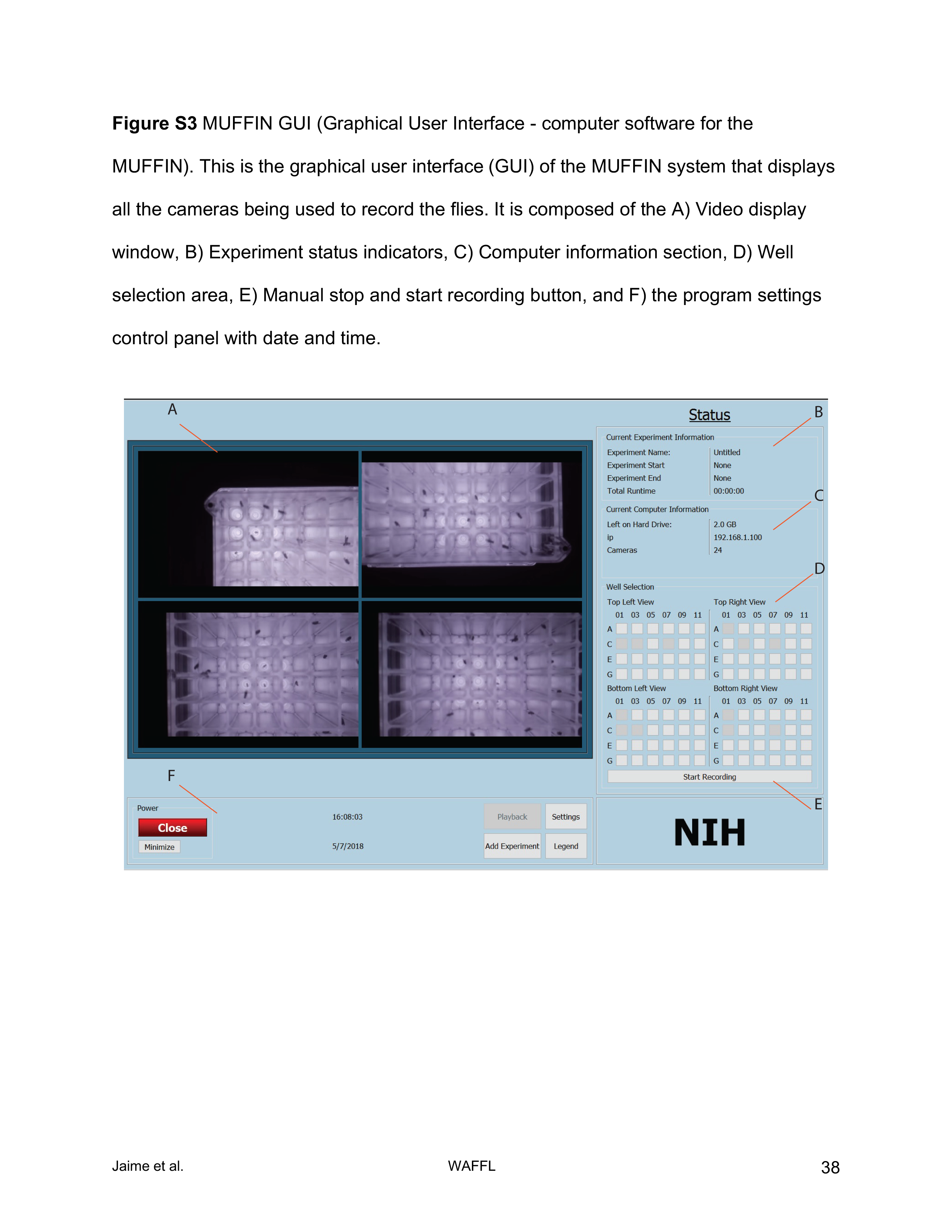
MUFFIN GUI (Graphical User Interface - computer software for the MUFFIN). This is the graphical user interface (GUI) of the MUFFIN system that displays all the cameras being used to record the flies. It is composed of the A) Video display window, B) Experiment status indicators, C) Computer information section, D) Well selection area, E) Manual stop and start recording button, and F) the program settings control panel with date and time.

